# A call for a unified and multimodal definition of cellular identity in the enteric nervous system

**DOI:** 10.1101/2024.01.15.575794

**Authors:** Homa Majd, Andrius Cesiulis, Ryan M Samuel, Mikayla N Richter, Nicholas Elder, Richard A Guyer, Marlene M. Hao, Lincon A. Stamp, Allan M Goldstein, Faranak Fattahi

## Abstract

The enteric nervous system (ENS) is a tantalizing frontier in neuroscience. With the recent emergence of single cell transcriptomic technologies, this rare and poorly understood tissue has begun to be better characterized in recent years. A precise functional mapping of enteric neuron diversity is critical for understanding ENS biology and enteric neuropathies. Nonetheless, this pursuit has faced considerable technical challenges. By leveraging different methods to compare available primary mouse and human ENS datasets, we underscore the urgent need for careful identity annotation, achieved through the harmonization and advancements of wet lab and computational techniques. We took different approaches including differential gene expression, module scoring, co-expression and correlation analysis, unbiased biological function hierarchical clustering, data integration and label transfer to compare and contrast functional annotations of several independently reported ENS datasets. These analyses highlight substantial discrepancies stemming from an overreliance on transcriptomics data without adequate validation in tissues. To achieve a comprehensive understanding of enteric neuron identity and their functional context, it is imperative to expand tissue sources and incorporate innovative technologies such as multiplexed imaging, electrophysiology, spatial transcriptomics, as well as comprehensive profiling of epigenome, proteome, and metabolome. Harnessing human pluripotent stem cell (hPSC) models provides unique opportunities for delineating lineage trees of the human ENS, and offers unparalleled advantages, including their scalability and compatibility with genetic manipulation and unbiased screens. We encourage a paradigm shift in our comprehension of cellular complexity and function in the ENS by calling for large-scale collaborative efforts and research investments.

## Enteric neurons are diverse

As the largest and most complex division of the autonomic nervous system, the enteric nervous system (ENS), holds the utmost significance in normal physiology and pathophysiology of the gut and disorders of gut-brain interaction (DGBIs). Comprising the cell bodies and projections of over 500 million neurons, and many more glia, the ENS is an intricate network that spans the length of the gastrointestinal (GI) tract. The ENS autonomously manages gut motility, secretion, absorption, local blood flow, and barrier function while also communicating with the brain, neuroendocrine and immune systems, and the microbiome. To fully understand the capabilities of the ENS, it is crucial to attain a comprehensive understanding of the molecular and functional diversity of its cell types. By unraveling the complex molecular and cellular processes, including epigenetic, transcriptional, post-transcriptional, translational, and post-translational regulatory mechanisms, as well as the context in which the ENS operates, such as the gut microenvironment, metabolic signals, and microbial and immune responses, we can begin to appreciate the remarkable autonomy and adaptability of the ENS in managing the diverse functions of the GI tract.

Towards this goal, ensuring accurate classification and characterization of enteric neurons is a critical first step with high-stakes implications for understanding the biology of the ENS and its associated diseases. Well-characterized cell populations and their markers serve as the basis for establishing *in vitro* and *in vivo* models, enabling researchers to explore ENS development, function, and crosstalk with other organ systems and external factors. Furthermore, consistent cellular annotations are crucial for scientific communication and interpretation of emerging research findings.

For decades, enteric neuron subtype identity has largely been defined based on morphological, immunohistochemical and functional features of neurons within the gut, including neuronal projection localization, neurochemistry, and electrophysiological properties. Investigations conducted on tissues isolated from model organisms, such as the guinea pig, mouse and rat^1^ have significantly progressed our understanding of the neurochemical and functional complexity of enteric neuron subtypes and have identified functional units within the ENS circuitry. These include intrinsic primary afferent neurons (IPANs or sensory neurons, SNs), ascending and descending interneurons (IN), excitatory and inhibitory motor neurons (EMNs and IMNs), and secretomotor and/or vasodilator neurons (SVNs)^1^. Despite the fact that many known neurotransmitters and neuropeptides are present in the ENS, only a handful are consistently used as markers to identify major cell types in the ENS. Some examples include nitric oxide synthase (NOS1) for IMNs and acetylcholine for EMNs as well as CALCA/B for IPANs^2^.

While it is impressive how large swaths of ENS function are coordinated by these neurotransmitters and neuropeptides, it also raises the question of whether yet-to-be-determined factors confer cell-type specific functional properties^3^. These features might include secreted peptides and proteins, electrophysiological features mediated by ion channels and transporters, metabolic profiles, and specialized capabilities for cell-cell interaction or synapse formation. To accurately understand the identity of individual cell types, it is also crucial to consider the contextual information in which they operate, including their position in the gut. Histochemical and functional assays have identified differences along the length of the GI tract, as well as between species, highlighting the need for more systematic efforts towards comprehensive mapping and profiling of the ENS^1^.

## Single cell transcriptomics identifies distinct enteric neuron subtypes

With the emergence of single cell and single nuclei transcriptomics (scRNA-seq and snRNA-seq) over the past decade, enteric neurobiologists have begun to overcome the gap posed by the limited availability of histochemical and functional probes to distinguish between subtypes of enteric neurons. In fact, several scRNA-seq datasets of the ENS have been recently published and reviewed, including mouse, human, and human pluripotent stem cell (hPSC) derived enteric neurons^4–12.The^ datasets of primary mouse and human intestine published by Ulrika Marklund (UM-mouse^8^, **Figure 1A**), Aviv Regev (AR-mouse and AR-human^4^, **Figure 1 B and C**), Sarah Teichmann (ST-human^9^, **Figure 1D**) and colleagues provide immensely valuable transcriptional profiling of a highly intricate biological tissue that is inherently challenging to acquire.

**Figure 1:**
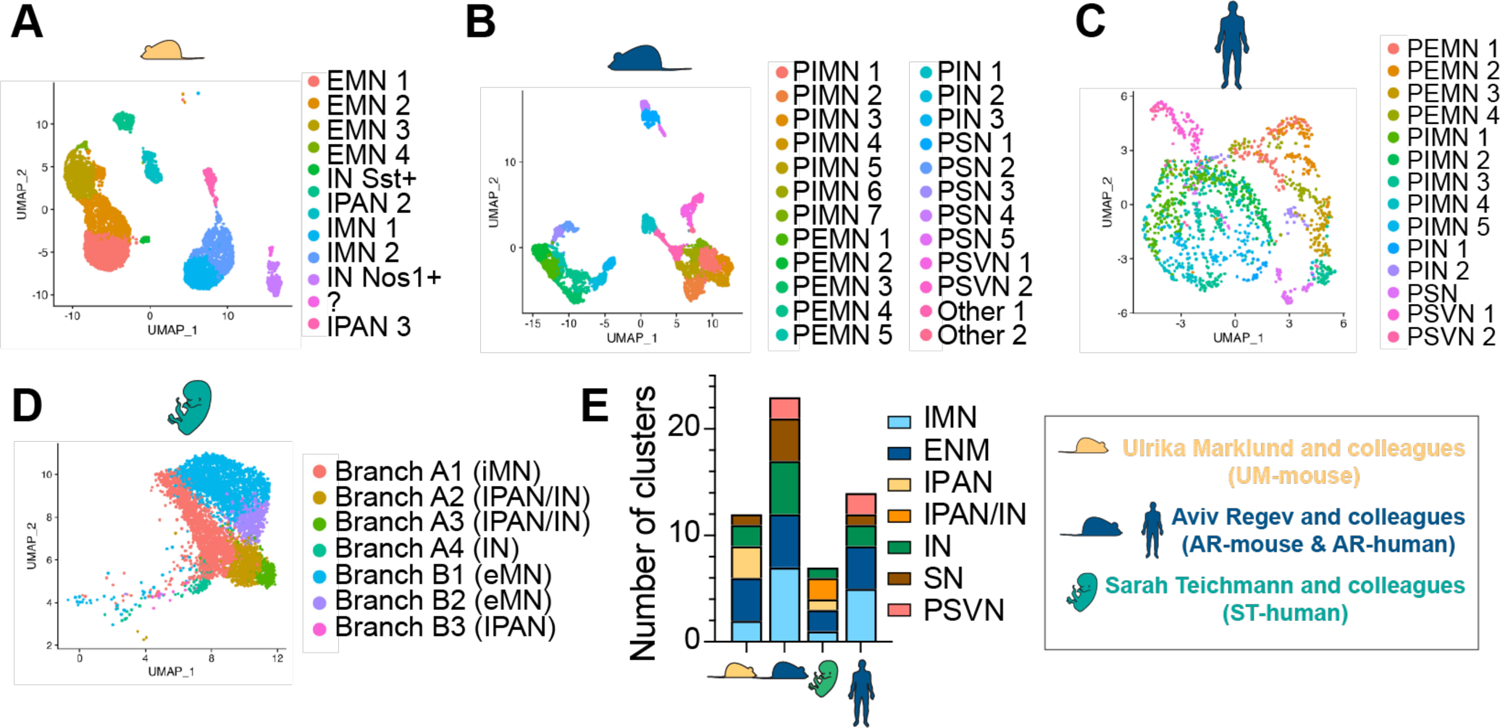
Primary mouse and human enteric neuron datasets used for cross dataset comparison **A-D**) UMAPs of enteric neurons generated from the original datasets of UM-mouse (**A**), AR-mouse (**B**), AR-human (**C**) and ST-human (D). “?” refers to the cluster labeled as “ENC11” or “?” in Morarach et al.^8^ **E**) Total number and distribution of enteric neuron cluster annotations **A-D**.

These studies present functional annotations for clusters of neurons, with varying numbers and identities of functional classes and subclasses (**Figure 1E**). For instance, the term IPAN is missing in annotations used in AR-human and AR-mouse, and PSN (putative sensory neurons) is used instead. Each dataset resolves different numbers of IMN clusters with UM-mouse containing two clusters, AR-mouse containing seven, ST-human containing only one, and AR-human containing five (**Figure 1E**). While these differences may also be due to subjectivity in clustering resolution and the analysis pipeline, the power of primary EN datasets has also been hindered by technical challenges arising from the limited number of cells in specific clusters, particularly in human studies. Furthermore, these datasets encompass primary ENS neurons from various species, distinct developmental stages, different regions of the GI tract, and employ different methodologies. This can be considered as both a strength and a caveat. On one hand, it provides valuable insights into the extent of neuronal diversity within the ENS. On the other hand, it presents challenges when attempting to infer the common features that define the identity of specific neuronal subtypes.

## Markers used for identification of neuronal subtypes in different datasets are inconsistent

Transcriptomics datasets of the ENS have been analyzed in isolation. Consequently, the list of genes used to annotate functional subtypes is different in each paper. Even among shared genes, the expression patterns vary widely across these datasets (Figure 2 A-P). We examined the complete list of genes used for annotating primary enteric neuron clusters, and found that only three, NOS1, TAC1, PENK were shared across the three studies (Figure 2A). Of the 32 markers in UM-mouse (Figure 2B), 5 (COX8C (human gene for mouse Cox8c), SLC18A3, NEUROD6, DLK1, DBH) were not detected in AR-human (Figure 2E). CALB, which is highly expressed in UM-mouse IPAN and EMN, was expressed highly in AR-human IMN and PSVN (Figure 2E). Another example is NTAFTC1, with a high expression in UM-mouse EMN and AR-human PIN (Figure 2E). The highest expression of GRP, a marker for ST-human IPAN/INs, is expressed by UM-mouse EMN 4 and AR-mouse PSNs (Figure 2 **F and H**). Transcripts of NEUROD6 and NXPH4, markers used by ST-human, were not detected in any AR-human clusters (Figure 2I). CHRM3, an IPAN marker in ST-human, is instead expressed by PSVN and PEMN clusters in the AR-human dataset (Figure 2I). SPOCK1, an IMN marker as defined by ST-human, was highly expressed by EMN 3, IPAN 2, IPAN 3, IN and SN in UM-mouse (Figure 2F), and by PEMNs, PINs, PSVNs, but not PIMNs, in AR-mouse (Figure 2G). Notably, even for the three shared genes across studies, cells positive for each marker were annotated differently (Figure 2 B-M **red boxes and** Figure 2 N-P). TAC1, a marker attributed to IPAN/IN in ST-human, is a PSN marker in AR-human and an EMN marker in UM-mouse.

**Figure 2:**
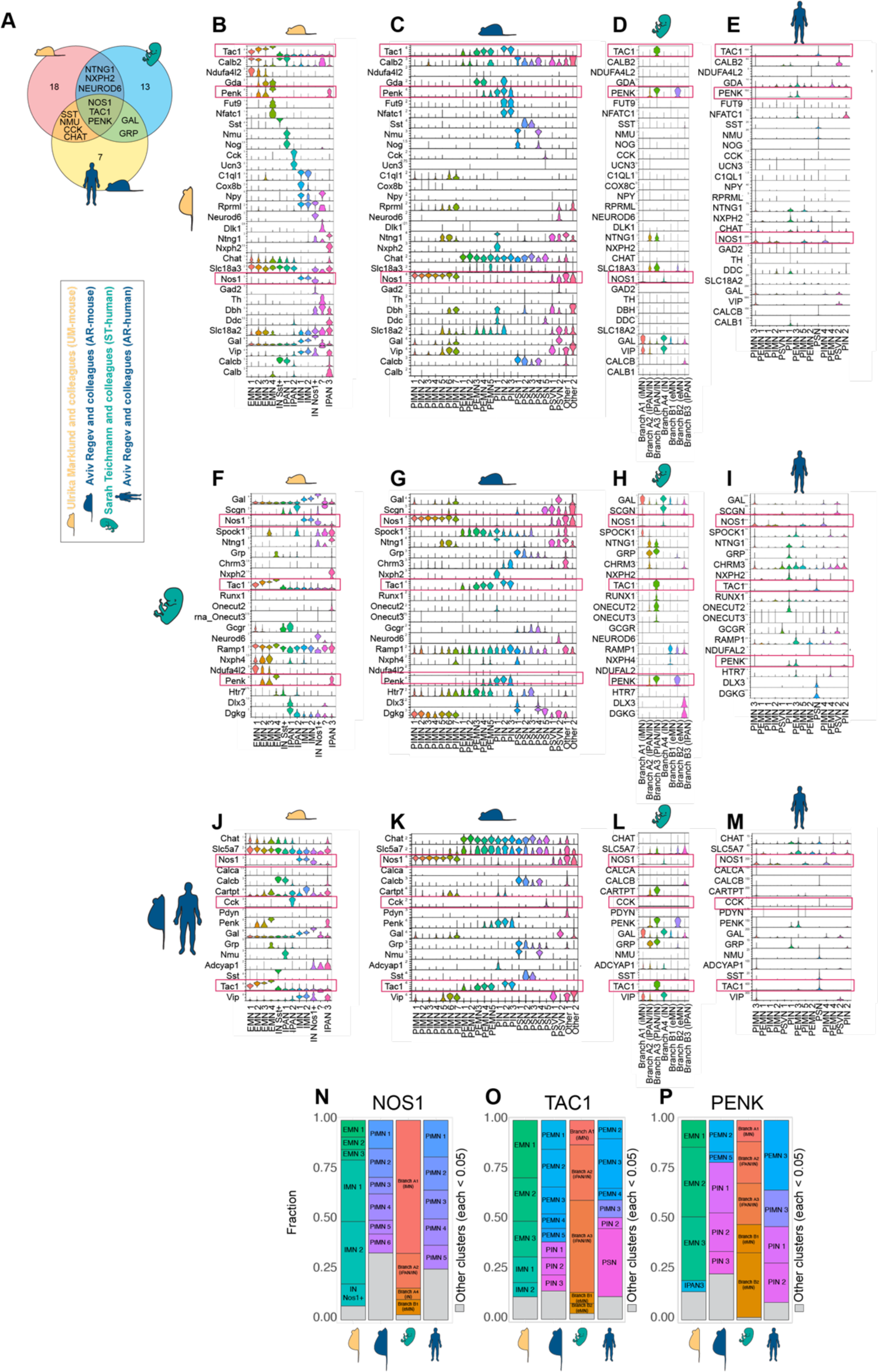
Cross dataset expression of primary enteric neuron cluster specific markers **A)** Venn diagram indicating shared markers used by each study. **B-E)** Violin plot stack of cluster specific markers originally used for UM-mouse, **B)** in UM-mouse, **C)** in AR-mouse, **D)** in ST-human, **E)** in AR-human. **F-I)** Violin plot stack of cluster specific markers originally used for ST-human, **F)** in UM-mouse, **G)** in AR-mouse, **H)** in ST-human, **I)** in AR-human. **J-M)** Violin plot stack of cluster specific markers originally used for AR-mouse and AR-human, **J)** in UM-mouse, **K)** in AR-mouse, **L)** in ST-human, **M)** in AR-human. *OR: **B-J)** Violin plot stack of cluster specific markers originally used for UM-mouse **(A-D)**, ST-human **(E-H)** ST-human and AR-mouse and human **(I-L)** all the datasets used in this analysis*. **N-P)** Comparison of the distribution of cluster annotations for enteric neurons expressing **(N)** Nos1 or NOS1, **(O)** Tac1 or TAC1, and **(P)** Penk1 or PENK1 transcripts across primary mouse and human ENS datasets used in this study.

Our systematic analysis of ENS datasets in parallel highlights substantial discrepancies between cluster-specific transcriptional markers, functional annotations, and the number of subtypes resolved in each dataset. These discrepancies suggest that much of the heterogeneity within the ENS remains to be determined and that annotating functionality based on the expression of only a handful of genes is simply insufficient.

## Beyond known markers: unbiased detection of neuronal subtypes across different datasets

Since the curated lists of identity genes used in various independent studies did not consistently mark enteric neuron subtypes, we reasoned that unbiased comparison of gene expression signatures could help identify shared and distinct cell populations. There are several categories of computational tools that enable such comparisons in scRNA-seq datasets^13^, including differential expression analysis, signature scores, co-expression network analysis, batch effect removal and data integration, annotation/label transfer, and trajectory analysis. Here, we employed some of these strategies to comprehensively compare these enteric neuron datasets.

## 1. Label transfer

Unbiased machine learning-based label transfer methods have emerged as valuable tools for annotating scRNA-seq and snRNA-seq datasets without the need for manual annotations^13,14^. Different methods have been developed which vary in their sensitivity to the input features, number of cells per population, and their performance across different annotation levels and datasets. These methods can be broadly categorized as supervised and unsupervised approaches, each offering distinct advantages and limitations. Supervised methods, including K-nearest neighbors (KNN) and Support Vector Machine (SVM), demonstrate robust performance when provided with a well-annotated and representative reference dataset. By leveraging gene expression similarities between cells in the reference data, these methods transfer cell type labels to query cells. However, supervised approaches may suffer from biases if the reference dataset does not match the heterogeneity of the query data. On the other hand, unsupervised label transfer methods can handle complex and heterogeneous cell populations without relying on pre-annotated reference data. These methods often learn intrinsic representations and identify cell clusters, enabling label transfer based on gene expression profile similarities. Nonetheless, they might encounter challenges when dealing with rare cell types and limited reference datasets. By rigorous comparison of various automated label transfer methods, previous studies have demonstrated that most classifiers perform well on a variety of datasets with decreased accuracy for complex datasets with overlapping classes or deep annotations^13,14^.

In our attempt to map neuronal subtypes across primary ENS datasets we used SingleCellNet^15^ (SCN), a random forest based supervised clustering algorithm, which has been shown to perform well when assessed for accuracy, percentage of unclassified cells, and computation time. Briefly, SCN operates by randomly selecting a sample from a reference dataset and training itself on the reference. The performance of the resulting algorithm is then evaluated based on how accurately it predicts the annotation of the remaining cells in the same dataset. We determined that using 100 cells was a suitable training set for the majority of clusters. When provided with a query dataset, SCN utilizes the reference training to annotate the query cells based on the reference annotations and assigns a ‘random’ identity when no reference annotation can be determined. We employed this approach to assess how the clusters of primary ENS neurons would have been annotated according to the other published ENS annotating criteria. Notably, this unbiased method relies solely on the transcriptional profile of each cell and is not biased by any prior annotations or assumptions. The abbreviations used here are consistent with the commonly used denotations in the field and in the original papers: IMN (inhibitory motor), EMN (excitatory motor), IN (inhibitory), IPAN (intrinsic primary afferent), PSVN (putative secretomotor/vasodilator) and SN (sensory). The inclusion of “P” in the AR-mouse and AR-human indicates “putative” as originally termed.

In our SCN comparisons (Figure 3 A-M), UM-mouse subtypes were best annotated when the SCN model was trained on the AR-mouse dataset (Figure 3B), with IMN 1 and IMN 2 predicted as PIMN 7 as well as UM-mouse EMN 1, 2 and 3 also predicted as AR-mouse PEMN 5, 5 and 4, respectively (Figure 3B). UM-mouse INs (IN Nos1+ and IN Sst+) were predicted to be AR-mouse PSVN2 and PSN (Figure 3B) and IPAN 3 was annotated as AR-mouse PIN 1. When UM-mouse clusters were annotated using ST-human as the reference, IPAN 3 matched as IPAN/IN (Branch A2) and IPAN 1 as IPAN (Branch B3), however, a high percentage of EMN 4 was predicted to be IPAN/IN (Figure 3F).

**Figure 3:**
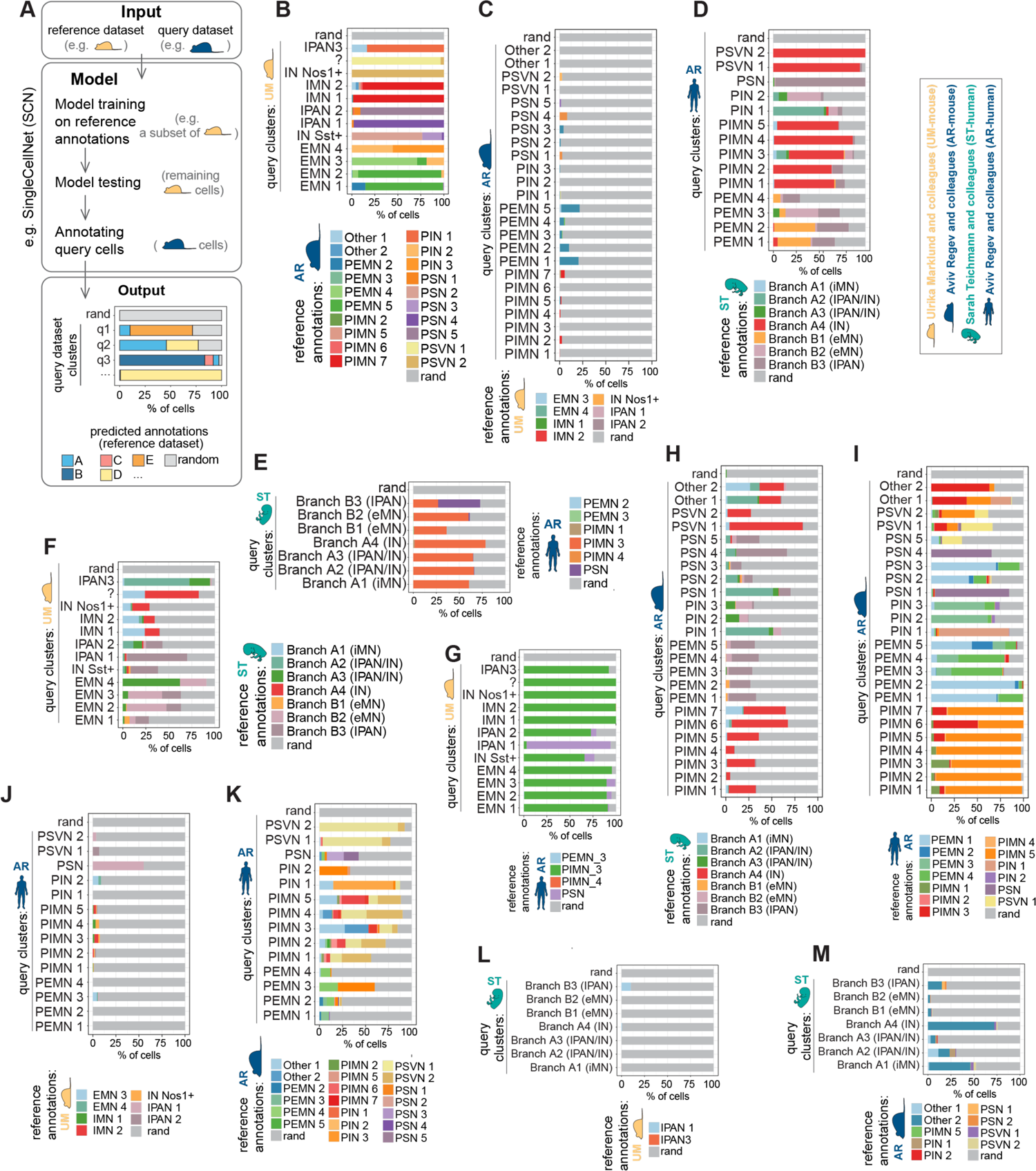
Unbiased cross dataset classification of primary enteric neurons using SingleCellNet **A)** Schematics of unbiased label transfer using SCN^15^. **B-M)** Reference primary enteric neuron scRNA-seq datasets of mouse (UM-mouse **(C, L, J)**, AR-mouse **(B, K, M)**) and human (ST-human **(D, F, H)**, AR-human **(E, G, I)**) were used to train SingleCellNet^15^. These models were then used for label transfer and cross annotation in the other datasets.

Taking AR-human as the reference dataset for annotating UM-mouse predicted PIMN identity for all the clusters except for IPAN1 which was predicted as PSN (Figure 3G). When the UM-mouse dataset was used as the reference, the model failed to predict the annotations that aligned with the functional neuron annotations employed in ST-human (Figure 3L), AR-human (Figure 3J) and even AR-mouse (Figure 3C) datasets. The majority of AR-mouse PEMNs were annotated as AR-human PEMNs with a predominant PEMN 1 and PEMN 4 (Figure 3I). For predicting AR-mouse identity, the AR-human annotations demonstrated the highest performance, which was to be anticipated given that both datasets originate from the same study (Figure 3I). All AR-mouse PIMN clusters (1-7) took the AR-human PIMN5 identity with only PIMN 6 also including a high proportion of PIMN 3 (Figure 3I). This suggests that the linear capture of biological diversity is not achieved either due to species differences or technical isolation challenges. PSN annotation was consistently captured for PSN 1 and 4 while PEMN identity was assigned to PSN 2 and PSN 3 as well as PIN 2 and PIN 3 (Figure 3I).

SCN analysis also revealed that human ENS neuron transcriptional profiles of neurons in clusters with the same functional annotations were not shared. AR-human PIMN annotation was assigned to cells from all of the clusters in ST-human query dataset (Figure 3E). According to ST-human as the reference, the annotation of AR-human PSVN 1 and 2 were almost entirely determined as IN (Figure 3D). ST-human IN identity was also predicted as the dominant annotation for AR-human PIMN 1-5 (Figure 3D).

Our unbiased supervised label transfer analyses further confirmed that annotations of neuron subtypes were specific to each dataset and not transferable across the board. These disparities raise intriguing questions regarding the extent to which they stem from limited statistical power, as opposed to meaningful biological differences. In addition, plasticity of the ENS is an important consideration that has been the topic of several recent studies. Alterations in gene expression patterns through the circadian cycle has been characterized in the murine ENS^4^, and changes in neuronal identify and function through the estrus cycle^16^. Overall, the findings underscore the importance of expanding tissue sources from diverse organisms, encompassing various developmental stages, and obtaining larger ENS datasets comprising a broader spectrum of neurons. Furthermore, the results highlight that rethinking annotation strategies, supported by rigorous experimental validations are essential to advance our understanding of enteric neuron identities and function.

## 2. Module scoring

As a common analysis method, module scoring is used to assess the activity or enrichment of pre-defined gene sets or modules in individual cells or clusters. By capturing coordinated changes in the expression of up to hundreds of genes within a module, module scoring can capture biologically significant relationships even if individual genes do not show strong differential expression. It also offers a tool to reduce the dimensionality of the data by summarizing the expression of gene sets in each cell, simplifying visualization and analysis. However, module scoring can be context dependent, and the relevance of predefined gene sets might be biased. Here, we used the top 100 differentially expressed genes for each cluster as the subtype signature modules in each dataset.

The analysis further reinforced the absence of consistent functional annotations for neuronal clusters based on transcriptional profiles (Figure 4 A-C). When the expression of ST-human and AR-human cluster modules was assessed in UM-mouse cluster, there was generally a consistent positive correlation between IMN clusters as well as between EMN clusters (Figure 4A**, blue boxes**). However, AR-human PIN 1 module was highly expressed by UM-mouse IPAN 3 and AR-human PIN 2 module was highly expressed by UM-mouse EMN 3 and EMN 4. The highest expression of AR-human PSVN 1 and PSVN 2 modules was observed in UM-mouse SN cluster (Figure 4A**, red boxes**). While AR-human PIMN 1-5 modules scored moderately positively in ST-human IMN cluster, their highest expressions were detected in Branch A4 (IN) for all except PIMN 3 which showed the highest expression in Branch A3 (IPAN/IN) (Figure 4B). Moreover, while AR-human PEMN 1-4 were positively scored in the two ST-human EMN clusters (B1 and B2), they were also scored positively in Branch B3 which is annotated as IPAN. In addition, AR-human PSVN 1 and PSVN 2 modules were highly expressed by ST-human Brach A4 (IN) cells (Figure 4B **red boxes**). When the expression of UM-mouse cluster modules was assessed in ST-human clusters, we again noticed unexpected correlations. For example, while UM-mouse IMN 1 and IMN 2 were positively correlated with ST-human IMN, they were much more highly correlated with ST-human Branch A4 (IN). As another example, UM-mouse EMN 2-4 modules were scored positively in ST-human Branch A3 (IPAN/IN) (Figure 4B**, red boxes**). Taken together, our module-based annotations indicate that the functional annotations of ENS neurons is subject to varying interpretation and accuracy when compared across different studies.

**Figure 4:**
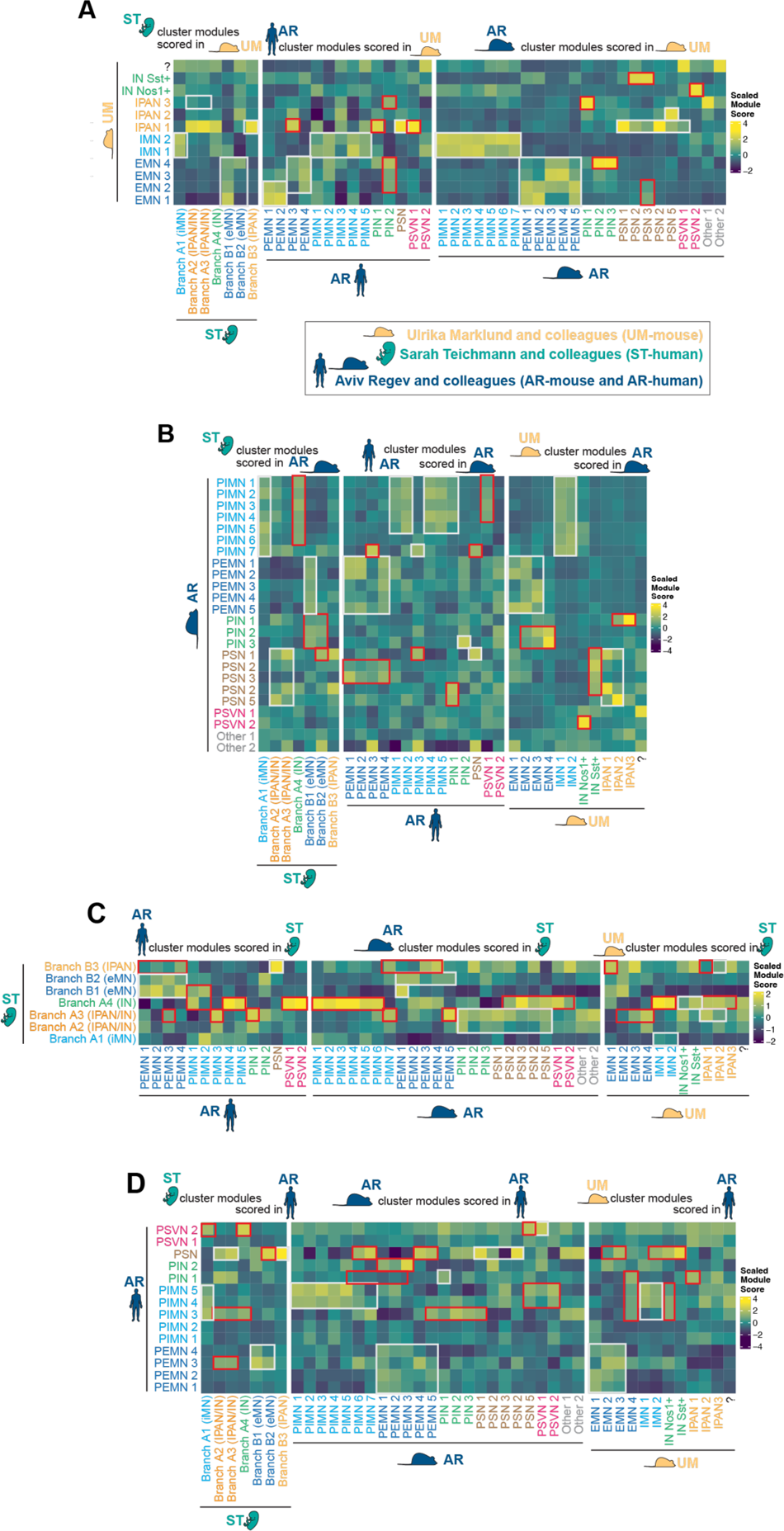
Cross-dataset module scoring for transcriptional signatures of primary enteric neuron clusters **A)** Heatmap of the average module scores of ST-human and AR-human neuronal subtype transcriptional signatures in UM-mouse. **B)** Heatmap of the ST-human, AR-human and UM-mouse neuronal subtype transcriptional signatures in AR-mouse. **C)** Heatmap of the average module scores of AR-human and UM-mouse neuronal subtype transcriptional signatures in ST-human. **D)** Heatmap of the average module scores of ST-human and UM-mouse neuronal subtype transcriptional signatures in AR-human.

## 3. Correlation analysis

Co-expression network analyses are computational methods used to identify and analyze co-regulated gene modules or clusters based on the patterns of gene expression across samples. As a co-expression network analysis, Spearman correlation analysis can be used to measure the similarity or dissimilarity of gene expression patterns between cell clusters or cell types identified in different scRNA-seq datasets. This allows researchers to explore the relationship and potential functional connections between cell populations across multiple datasets. Some of the key advantages of Spearman correlation compared to other co-expression network analysis (such as Pearson correlation) include its robustness to outliers but, similar to other computational methods, it is sensitive to small sample size. A high positive Spearman correlation between the gene expression patterns of two clusters suggests functional similarity. However, a negative Spearman correlation indicates distinct gene expression patterns and, therefore, potential functional differences. A Spearman correlation close to 0 suggests little to no relationship between clusters.

To compare enteric neuron subtypes in different datasets, we performed Spearman correlation analysis using 3000 and 100 anchor features. Even though the 100-feature analysis showed higher similarity scores overall, the majority of functional annotations did not align between the clusters of different studies (Figure 5 **A and B**). Overall IMN and EMN clusters showed positive correlation between datasets with the mouse datasets exhibiting stronger correlation compared to the human datasets, likely due to more accurate annotations as a result of more extensive functional and molecular data in mice. However, the relationship between motor neuron subtypes was not a one-to-one correlation, as anticipated due to the presence of varying subtypes within each motor subtype. Furthermore, both the human-human and mouse-mouse analyses revealed clear discrepancies between the datasets, with Figure 5 **A and B** highlighting consistent (blue boxes) and inconsistent (red boxes) functional annotations. For example, the highest correlation score for the ST-human IN cluster was with AR-human PIMN4 while correlation with AR-human PIN clusters was negative (Figure 5A). While the highest correlation of UM-mouse EMN1-3 (.91, 0.8, .86 respectively) was scored in AR-mouse PEMNs, the highest correlation score of UM-mouse EMN4 was with AR-mouse PIN 2 and 3 (.86, 0.83 respectively) (Figure 5B).

**Figure 5:**
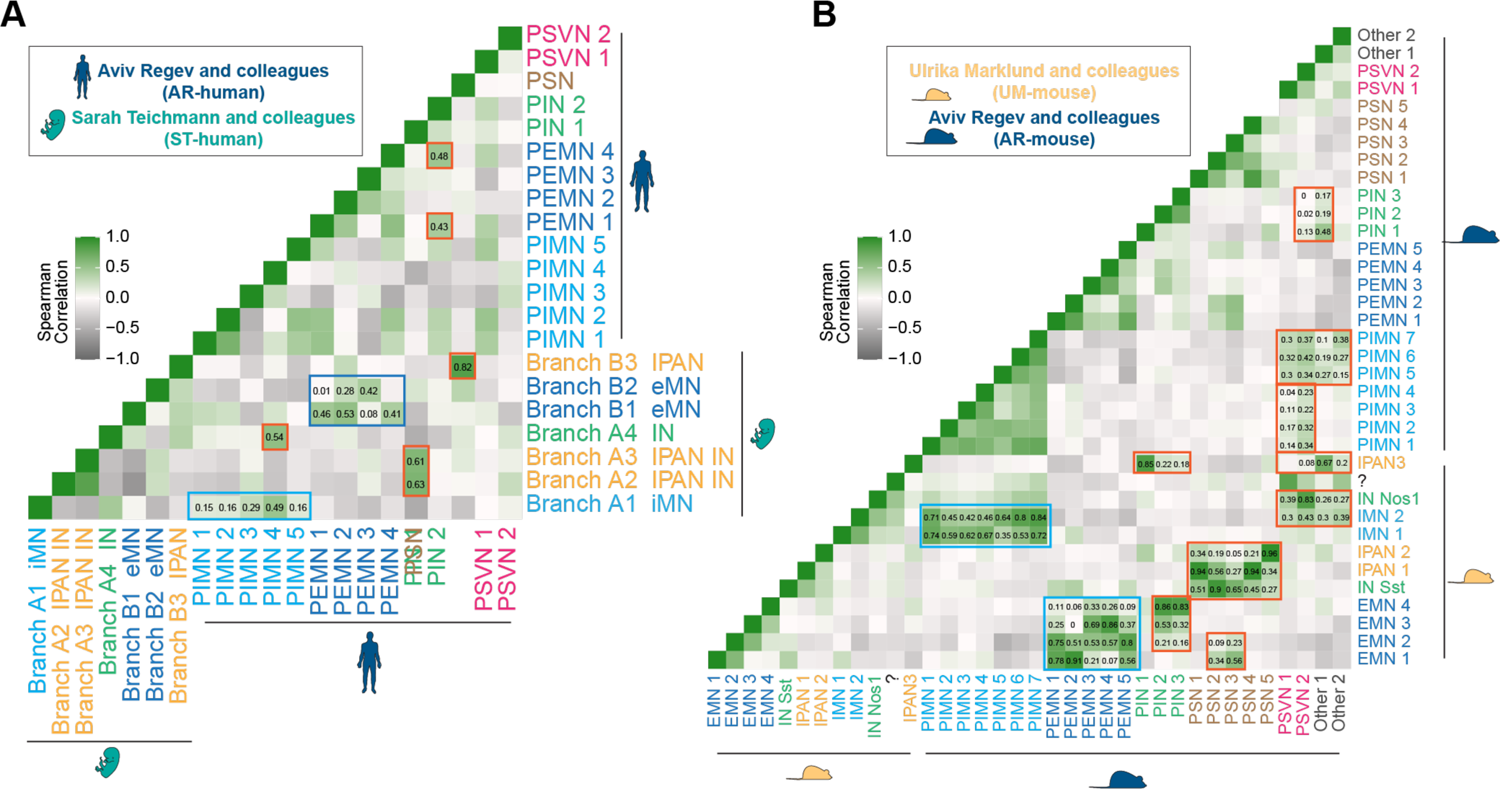
Comparative analysis of primary ENS neurons using Spearman correlation analysis **A-B)** Heatmap matrix of Spearman correlations based on expression of 100 anchor features shared significantly variable genes (or anchor features) between **A)** primary human (ST-human and AR-human) and **B)** primary mouse (UM-mouse AR-mouse) enteric neuron subtypes.

Our correlation analysis of enteric neuron datasets uncovers substantial gene expression differences between clusters that are annotated similarly in different datasets. While some positive correlations exist for well-characterized functional subtypes (such as some motor neuron populations), the overall similarities remain modest. These disparities again highlight the insufficiency of transcriptional comparisons to assign functional annotations and the potential revelation of the ENS being more complex and possibly more functionally diverse than previously identified. Thus the collection and analysis of additional and larger ENS datasets backed by rigorous functional experimental validations is warranted.

## 4. Integration

Data integration methods aim to merge multiple datasets to create a unified representation of cell types^17^. Various integration methods have been developed to align and harmonize datasets into a shared low-dimensional space, enabling comparison across datasets. By incorporating noise reduction and batch correction techniques, integration strategies aim to retain biologically relevant information and preserve the underlying biological relationships between cells. Cells that represent similar gene expression profiles across different datasets tend to group together and enable more accurate downstream analysis such as clustering and differential expression analysis as well as label transfer. Harmony^18^ is a graph-based integration method that aims to capture the similarities and distances in gene expression profiles of different datasets. The integrated graph resulting from aligning the graph structures of different datasets enables direct comparison and visualization.

Integration of UM-mouse and AR-mouse ENS datasets shows that while some of the Harmony clusters in the integrated UMAP space are composed of cells from both datasets, five of the 14 clusters are almost entirely derived from one dataset and six other clusters are >75% composed of cells from one dataset (Figure 6 A-C). Cluster 6 is almost 50% derived from UM-mouse IPAN and 50% AR-mouse PIN. Clusters 7 and 14 show a balanced integration of UM-mouse IPAN and AR-mouse PSN (Figure 6 A-C). So, while two clusters appear to properly integrate cells with similar functional annotations, this is not the case for the rest of the integrated dataset and the datasets largely remain distinct post-integration.

**Figure 6:**
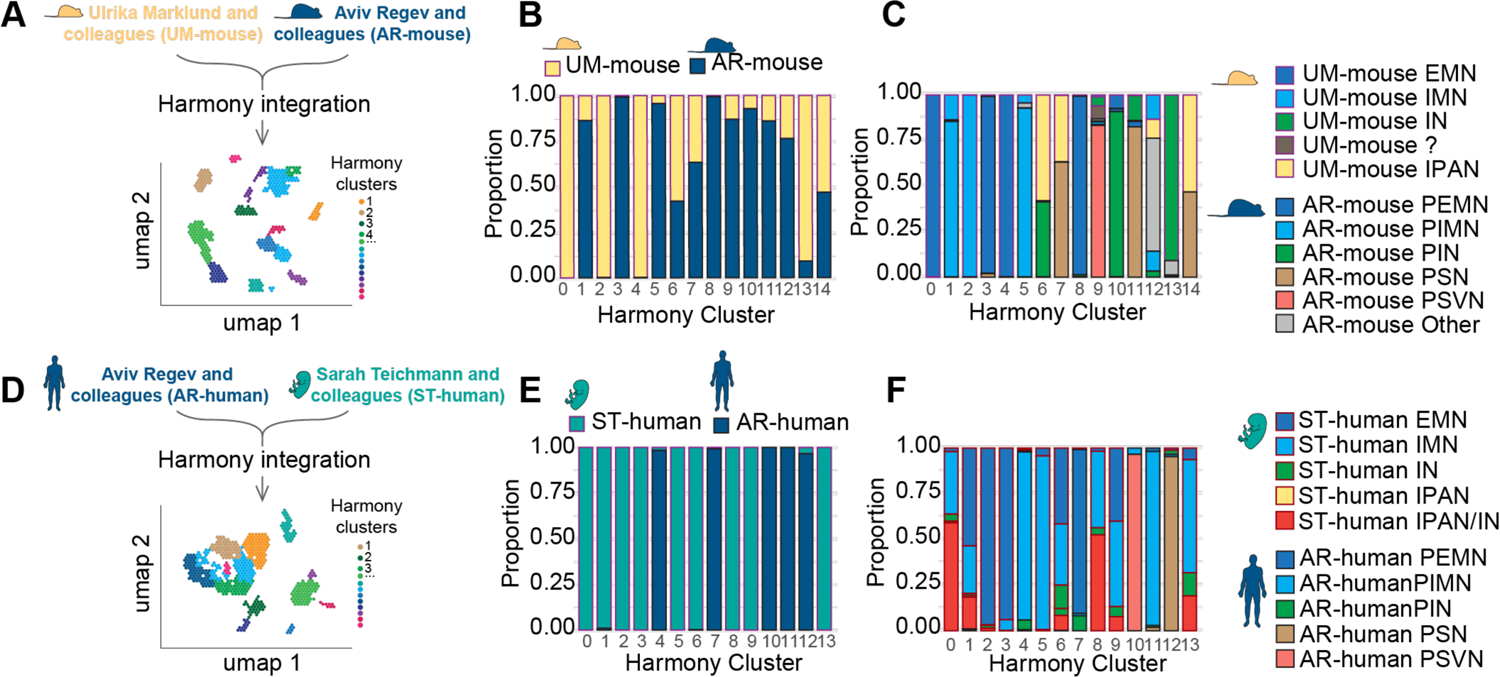
Harmony integration of primary mouse and human enteric neuron datasets. **A)** Schematic representation of Harmony integration of UM-mouse and AR-mouse datasets. **B, C**) Distribution of cells derived from UM-mouse and AR-mouse datasets (**B**) and their respective broad functional annotations in each Harmony cluster (**C**). **D**) Schematic representation of Harmony integration of ST-human and AR-human datasets. **E, F**) Distribution of cells derived from ST-human and AR-human datasets (**E**) and their respective broad functional annotations in each Harmony cluster (**F**).

The integration of ST-human and AR-human datasets (Figure 6D) revealed distinct differences in the transcription profiles of cells from each dataset. Each of the 13 Harmony clusters were almost entirely derived exclusively from either ST-human or the AR-human datasets (Figure 6 **E and F**). While this clear divide in integration might be attributed to variations in the developmental stages of the neurons, the absence of any integration, even in the well-studied motor neuron clusters, underscores the need for a critical review and comprehensive analysis of the annotations, sample preparation, and development of improved computational and analysis techniques.

Altogether, to gain a deeper understanding of the observed discrepancies and ensure accurate cluster annotations, it is essential to thoroughly assess the experimental procedures, data processing pipelines, and potential batch effects. By addressing these challenges, we can enhance the reliability and validity of the integration results and advance our understanding of neuronal diversity within the ENS.

## 5. Clustering based on gene set enrichment analysis (GSEA)

Hierarchical clustering is a valuable unsupervised clustering strategy to help identify similarities and differences in gene expression patterns among cells or clusters of different datasets. It offers a particularly useful method when assessing the overall relationship and similarities between different populations across multiple datasets, identifying conserved cell types, as well as identifying potentially novel cell populations without biasing the analysis with prior knowledge. It constructs a tree-like structure (dendrogram) where branches are formed through merging and splitting based on similarity. Like other methods, it is essential to consider batch effects, data normalization, and sample size. Other unbiased clustering methods include K-means clustering which requires the user to specify the K number of clusters in advance, or graph-based clustering which leverages the graph structures and identifies clusters based on their relative similarities compared to other cells/clusters.

To assess the functional similarities of primary mouse and human ENS neuron subtypes based on defined gene sets, we conducted gene set enrichment analysis (GSEA) using the gene ontology biological process (GOBP) gene sets. Comparing clusters based on the enrichment of a large number of gene sets offers solutions to many technical challenges when comparing independently produced datasets. Attributing the expression of numerous genes to a single GO pathway allows for the detection of common pathways, even if different subsets of the gene set were detected in different datasets. This overcomes technical limitations attributed to batch effects and gene dropout due to differences in sequencing depths between datasets. Additionally, this method allows for cross species comparison without the need for homologous gene assumptions, as the same pathways have been annotated for both mouse and human specific genes and gene family members.

We performed this analysis on the significantly upregulated gene lists of each neuron cluster, which were calculated independently for each dataset and rank ordered by log2 fold change. We employed hierarchical clustering across all datasets, using the normalized enrichment score of enriched GO biological process terms found in at least one cluster (Figure 7). This clustering revealed only six instances of close relationships among similarly annotated neuron clusters from different datasets. Notably AR-mouse PSN 1 and 4 cluster with UM-mouse IPAN 1 and ST-human Branch B3 IPAN, respectively, while the remaining instances consist of groups of EMNs or IMNs from all four datasets. However, the majority of the dendrogram shows close relationships of clusters with diverging functional annotation both across and within datasets. For instance, AR-human PIN1 clusters closely with UM-mouse EMN4, and AR-human PIMN1, PIN 2 and PSVN 1 all are predicted to be closely related. Overall, these results suggest that while many of the primary enteric motor neurons (IMNs and EMNs) exhibit close clustering, there are notable discrepancies in the overall functional clustering and/or annotation of neuronal subtypes (Figure 7). A similar analysis performing hierarchical clustering based on all GO biological process, molecular function, cellular component and human phenotype pathways showed similar results to the biological process pathways alone (data not shown).

**Figure 7:**
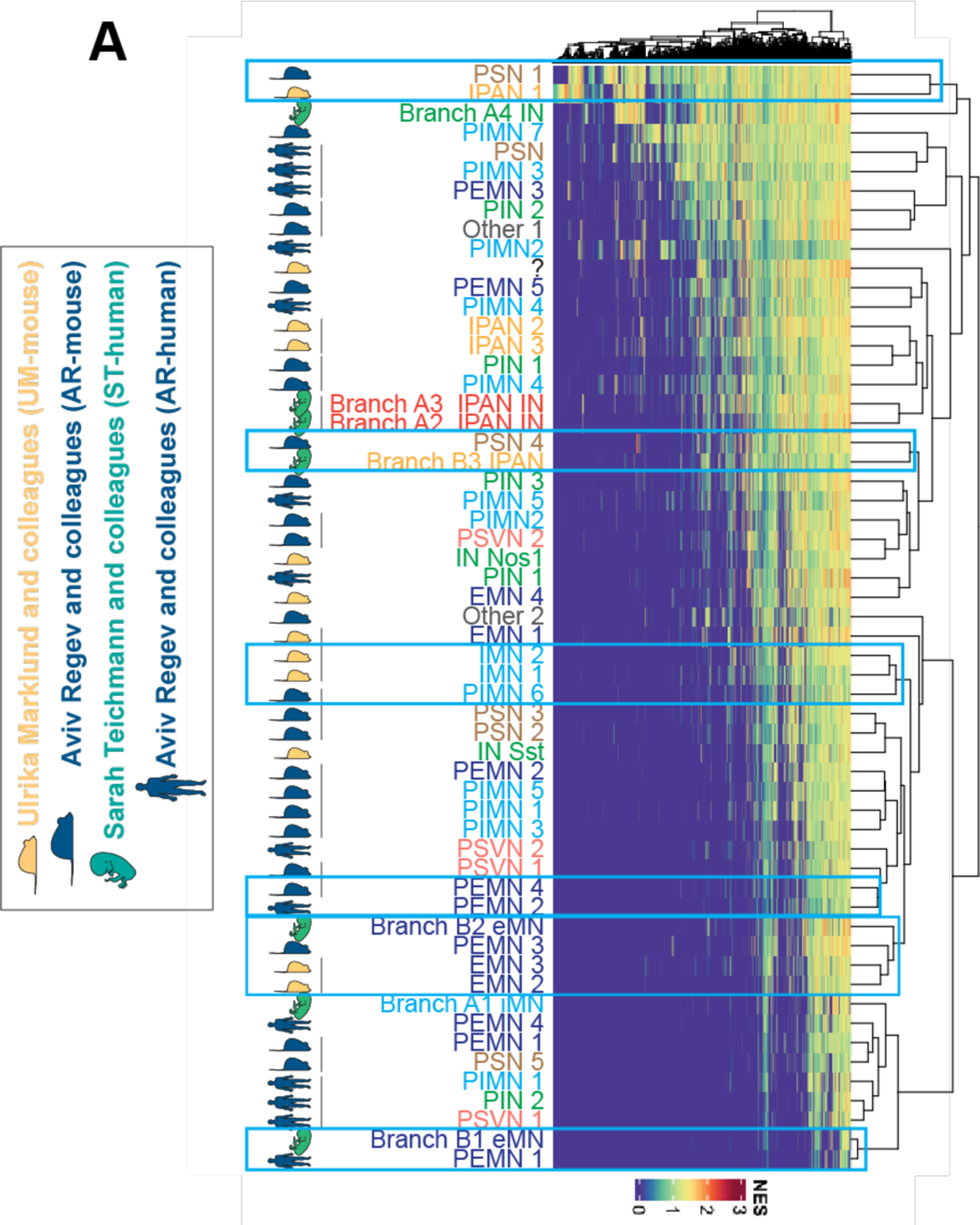
Comparative analysis of primary ENS neurons using GOBP hierarchical clustering. Hierarchical clustering of primary enteric neuron clusters based on normalized enrichment scores of biological process gene ontology (GOBP) pathways. Blue boxes indicate closely clustered neuronal subtypes with matching functional annotation from two different datasets.

The unbiased clustering based on enrichment scores of GOBP reveals a dendrogram consisting of closely related clusters with different functional annotations. This suggests the presence of functional differences derived from the transcriptome profiles that were not adequately captured during the initial cluster annotations using only a few markers. The diverse annotations within seemingly similar clusters emphasize the necessity for a more comprehensive and thorough examination of the data when annotating clusters, even when solely considering the transcriptome as a readout of cellular identity. It is, however, important to note that this analysis and these observations are constrained by the availability of well-defined pathways related to our cell types of interest. To date, many GO biological process pathways have been describe for various neuronal functions, such as synaptic transmission and response to various neurotransmitters. However, the full breadth of functions and processes performed by enteric neurons has yet to be described and mechanistically dissected to allow for their inclusion in the above analysis.

## Conclusions

Transcriptomic datasets of the ENS from various species, sexes, age groups, GI regions, gut layers, and isolation techniques have provided valuable resources for researchers in the field. However, it is important to acknowledge that these factors contribute to inherent differences that must be taken into consideration before attempting to generalize findings. Furthermore, our knowledge of these differences, even within a single organ region, organism, or tissue layer, is limited compared to what we know about the central nervous system (CNS). Thus, it is crucial to exercise caution in defining subtype specific markers used for annotation. Paradoxically, reaching clear conclusions from ENS transcriptomics are becoming more challenging as more data is generated. We believe this largely stems from the lack of a unified system for classifying cell identities, lack of clear criteria for defining distinct categories, and overreliance on transcriptomic. Because many post-transcriptional processes impact protein expression, RNA transcripts are only imperfectly correlated with phenotypes. The extent to which transcriptional differences in individual neurons indicate functional differences is not clear, and should be a high priority area for investigation.

Another layer of complexity in our understanding of the ENS is its cellular plasticity. In contrast to the CNS, where cell fate is largely determined and differentiation is set early in development, the ENS retains a remarkable degree of plasticity throughout an organism’s life ^19–22^. ENS cells, derived from neural crest origins, are involved in an ongoing process of migration, proliferation, and maturation, accommodating the ever-changing landscape of the gastrointestinal tract. This dynamic capacity allows the ENS to adjust to variations in gut size, functionality, and to a range of environmental factors, including dietary shifts, inflammatory states, and injury. Neurons may modify their transcriptional profiles and change their neurotransmitter or receptor expression in response to environmental stimuli, similar to the transient and adaptive responses observed in enteric glia^23,24^. Consequently, the assumption that functionally distinct neurons equate to permanent cell states, or lineages may be overly simplistic and may not account for the nuanced nature of neuronal identity. The full extent of this dynamic nature of the ENS is still not well understood, which further complicates the annotation, clustering, and subsequent analysis of these cells.

While transcription profiling yields critical insights into cellular functions, it represents just one dimension of the complex regulatory network that dictates gene expression and cellular identity. Beyond transcriptional regulation, there are multiple layers of post-transcriptional control—such as RNA stability, processing, editing, transport, and alternative splicing—that greatly influence the final functional outcomes. These mechanisms enrich the diversity of the proteome but are often not fully detected by short-read RNAseq, necessitating long-read sequencing techniques to reveal such RNA variants. Furthermore, the translation of mRNA into protein is governed by factors including ribosome recruitment, initiation, and elongation rates, as well as regulatory sequences within the mRNA itself. Post-translational modifications (PTMs) like phosphorylation, acetylation, methylation, ubiquitination, among others, critically modulate protein functionality, stability, and cellular location, impacting protein activity and signaling pathways. Therefore, transcriptomic clusters may not accurately reflect the functional diversity of neuronal populations and mathematical clustering may identify distinct groups that do not necessarily correlate with significant functional differences (Figure 8). Careful verification and thorough characterization of identified transcriptomics markers, exemplified by the elegant histological validation using primary mouse tissue in the study by Morarach et al.^8^, provide a robust template for conducting validation studies of this nature.

**Figure 8:**
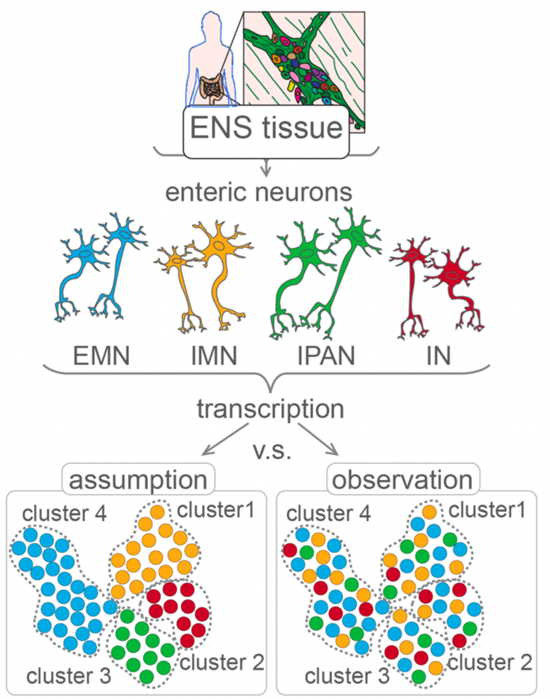
Transcriptional identities are not synonymous with functional identities in enteric neurons

Accurate characterization and appropriate annotation of enteric neuron functions require moving beyond relying on a single biological modality, such as transcription (Figure 9). By adopting a multidimensional approach, we can go beyond our current understanding of enteric neuron identity and unravel the complexities of the ENS.

**Figure 9:**
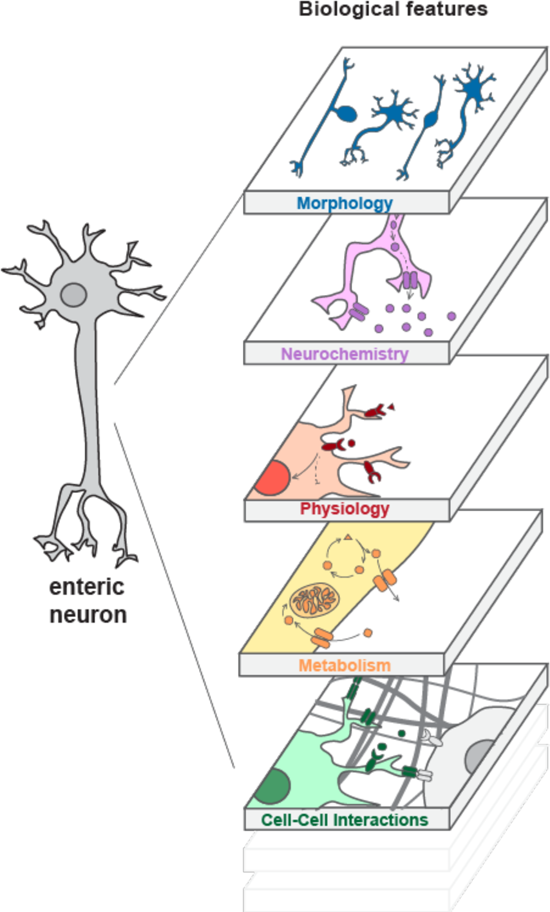
Enteric neuron identity should be defined based on multiple biological features

These profiling studies should be performed on diverse ENS samples that encompass a broader range of gut regions, animal models, developmental and disease states, gender and ethnic diversities ^6,12^. In parallel, the development of improved computational methods, including dataset integrations and label transfer algorithms, can facilitate the identification and classification of distinct ENS cell types and subtypes. These computational approaches can help uncover the shared and unique features of enteric neurons obtained from various sources. Moreover, integration and side-by-side analysis of tissue datasets from different sources, particularly human patients, requires caution due to potential biological and pathological differences, cell type variations, and diverse cell identities. Overlooking these factors can result in misleading analyses and inaccurate conclusions. In addition, the incorporation of spatial profiling and advanced imaging techniques, such as tissue clearing and organ-level imaging, holds promise for elucidating the spatial organization and connectivity of enteric neurons within the gut. It will ultimately be essential to unify the transcriptomic identities with the functional roles of enteric neurons. One route to this goal would be Patch-seq, which combines whole-cell patch clamp electrophysiological recordings and pharmacological analysis, with transcriptomic analysis^25^. Here, following electrophysiological recording of myenteric and submucosal neurons, the patch electrode would be used to collect the nucleus from the recorded cell for downstream snRNA-seq. Dye labelling of the recorded cell would also enable post-hoc tracing of its projections. While these would be technically challenging and low-throughput studies, together with spatial and imaging techniques, these approaches can reveal the regional diversity and spatial relationships of ENS cells, enhancing our comprehension of their functional roles.

Recently, it has been argued that a nomenclature annotation rooted in a consensus ontogeny, which encompasses the developmental history of cells throughout the life cycle of a specific species, would be preferable to using an atlas to establish a reference taxonomy for cell types^26^. These strategies aim to integrate lineage histories and molecular states across different species, which is an ideal approach. Understanding the developmental decisions of enteric neurons is essential. Techniques such as lineage tracing and multi-omics approaches can provide insights into the developmental trajectories and fate decisions of enteric neurons, enabling a better understanding of their functional diversity as well as ENS developmental neuropathies. The increasing accessibility of primary ENS datasets from mice and other organisms makes this approach more practical and valuable. However, studying the developmental trajectories of the human ENS using these strategies remains challenging due to tissue accessibility limitations and low yield with current isolation techniques. Considering this, the development of robust and reliable human pluripotent stem cell (hPSC)-based enteric neuron cultures can overcome these challenges and enable developmental studies, including lineage tracing experiments. Although hPSCs models do not fully replicate higher-level cell-cell interactions and primary ENS environmental cues, these cultures offer scalable and diverse sources of enteric neurons that can be obtained at various stages of differentiation. Furthermore, they are compatible with genetic manipulation and high-throughput screens. This allows for high-resolution lineage barcoding and tracing experiments, which can uncover the mechanisms of cell type specification in the human ENS. As a result, they serve as a valuable complement to current in vivo models ^7,27,28^.

In summary, the comparisons of existing primary enteric neuron datasets highlight the importance of recognizing and embracing the complexity of the ENS, urgent need for more cautious annotation, and the necessity to employ rigorous approaches to ensure robust data that will facilitate breakthroughs in the field. Gathering a consensus on transcriptomic and functional data on the ENS will enable better understanding of how deviations can culminate in disorders of gut-brain interaction (DGBIs), where the cause of gut dysfunction currently remains largely elusive. As ENS research continues to evolve, ongoing efforts to enhance the experimental and computational methods and address their limitations will likely lead to more accurate and comprehensive cell type identifications. By defining enteric neuron functional identities, these endeavors will open new avenues for transformative discoveries.

## Methods

### Spearman correlations

The transcriptional correlation of between cell clusters in the two datasets was computed using non-imputed gene counts and Seurat’s integration functions to first find 100 anchor features based on the first 30 dimensions of the canonical correlation analysis and then integrate the two datasets using the same number of dimensions. The expression of these 100 anchor features was then scaled and centered in the merged data object and the average scaled expression of each anchor feature was calculated for each dataset’s cell clusters of interest using the “AverageExpression” function. A Spearman correlation matrix comparing all cell clusters to all cell clusters was generated based on the average scaled expression of the 100 anchor features.

### Cell type transcriptional signature module scoring

To find transcriptionally similar neuronal subtypes between two datasets, first the differentially expressed (DE) genes of the reference dataset are calculated from the non-imputed gene counts with the “FindAllMarkers” function using the Wilcoxon Rank Sum test and only genes with a positive fold change were returned. The DE gene lists are first filtered to remove genes not present in the query dataset. Then for each cell cluster in the reference dataset, a transcriptional signature gene list is made from the top 100 (or as many as possible if <100) DE genes sorted first by LogFC and then increasing adjusted p-value (if two genes has the same LogFC). The query dataset is then scored for the transcriptional signature gene lists of each reference dataset cell cluster using the “AddModuleScore” function based on the query dataset’s gene counts (RNA assay, non imputed).

### Label transfer

We used SingleCellNet (SCN)^15^ for unbiased classification of primary enteric neuron subtypes. Reference datasets included studies published by Ulrika Marklund (UM-mouse^8^), Aviv Regev (AR-mouse and AR-human^4^), Sarah Teichmann (ST-human^9^) and colleagues. Feature expression matrices and associated metadata objects were derived from each dataset and used as reference or query datasets. For classification, model training custom parameters were used for each reference dataset. Model was trained on a subset of 100 randomly selected cells for each cell cluster present in the REF reference dataset, selecting top 20 DE genes and top 50 gene pairs for training.

### Harmony integration

Harmony^18^ integration was performed using the Seurat v5 harmony wrapper function. Enteric neuron datasets of the same species were first merged, and metadata was added to track each datasets equivalent of a unique biological sample (termed Dataset_SampleID) by appending a dataset identifier to the existing sample identifier column specific to each dataset (Sample.name for ST-human, Patient_ID for AR-human, Biorep for UM-mouse and Mouse_ID for AR-mouse). Due to the subsetting of each dataset to only include the original author annotated neurons, some samples contributed very few cells to the neuron subset dataset. Samples that consisted of less than 10 cells were removed to comply with the functionality of the subsequent data processing steps. First, the RNA assay was split to create a separate layer for each Dataset_SampleID. Counts normalization and variable feature identification is then performed on each layer separately to identify a consensus set of variable features across all samples. These shared variable features are then scaled and used for principal component analysis. The individual samples were then integrated using the “IntegrateLayers” function with the method set to HarmonyIntegration. The nearest neighbor graph construction and UMAP dimensionality reduction were then performed on the first 30 harmony components. Clusters were identified using the Louvain algorithm with a resolution of .3 and .6 for the integrated mouse and human datasets, respectively.

### GSEA hierarchical clustering

For each primary ENS dataset, DE genes for each For each primary EN dataset, DE genes for each neuronal cluster were calculated using the “FindAllMarkers” function. Gene set enrichment analysis (GSEA) was performed on each neuronal subytpe’s upregulated DE genes (positive log2 fold change only) sorted by decreasing log2 fold change using fgsea v1.16 for the MSigDB gene ontology (C5) pathways. Normalized Enrichment Scores (NES) were calculated for gene sets containing a minimum of 15 genes in the DE gene list with the scoreType set to “positive”. Recovered enriched pathways were filtered to only include biological process (GOBP) gene sets but not filtered based on significance as to not limit the result to pathways enriched in the highest fold change genes. The NESs of the filtered GSEA results for all clusters were then merged and pathways not detected in a neuronal cluster were assigned a NES of 0. Euclidean distance based hierarchical clustering was then performed based on the NESs to cluster both the gene ontology pathways and the neuronal clusters.

## Acknowledgements

The work was supported by grants from UCSF Program for Breakthrough Biomedical Research and Sandler Foundation, the NIH Director’s New Innovator Award (DP2NS116769) to F.F. and the National Institute of Diabetes and Digestive and Kidney Diseases (R01DK121169) to F.F., (F32DK121440) to R.A.G., and (R01DK119210) to A.M.G. H.M. is supported by Larry L. Hillblom Foundation postdoctoral fellowship, NIH T32-DK007418 fellowship and UCSF Program for Breakthrough Biomedical Research independent postdoctoral fellowship. We are grateful to the members of Fattahi Lab at UCSF for their constructive feedback on the manuscript.

## Author contributions

H.M. Design and computational analysis, writing of manuscript. A.C. computational analysis, writing manuscript. R.M.S. computational analysis, writing of manuscript. M.N.R. writing of manuscript. N.E. writing of manuscript. R.A.G. writing of manuscript. A.M.G. writing of manuscript. F.F. Deign and conception of the study, supervision of all efforts, writing of manuscript.

## Declaration of interests

F.F. is an inventor of several patent applications owned by UCSF and MSKCC and Weill Cornell Medicine related to hPSC-differentiation technologies including technologies for derivation of enteric neurons and their application for drug discovery.

---

**Table S1:** SCN assessment parameters.

| reference-for-query | kappa | AUPRC_w | AUPRC_wc |
| --- | --- | --- | --- |
| ST-human for mouse | 0.739 | 0.813 | 0.893 |
| ST-human for human | 0.737 | 0.827 | 0.893 |
| AR-human for mouse | 0.776 | 0.875 | 0.871 |
| AR-human for human | 0.779 | 0.876 | 0.881 |
| UM-mouse for mouse | 0.900 | 0.929 | 0.967 |
| UM-mouse for human | 0.875 | 0.913 | 0.948 |
| AR-mouse for mouse | 0.862 | 0.893 | 0.942 |
| AR-mouse for human | 0.861 | 0.915 | 0.942 |

## Notes

### Summary of Updates

the revised version of the manuscript includes corrections to mistakes previously identified in the list of references.

